# Transgenic microalgae expressing double-stranded RNA as potential feed supplements for controlling white spot syndrome in shrimp aquaculture

**DOI:** 10.1101/2022.02.09.479659

**Authors:** Patai Charoonnart, Henry N Taunt, Luyao Yang, Conner Webb, Saul Purton, Colin Robinson, Vanvimon Saksmerprome

## Abstract

Viral infection of farmed fish and shellfish represents a major issue within the aquaculture industry. One potential control strategy involves RNA interference of viral gene expression through the oral delivery of specific double-stranded RNA (dsRNA). In previous work we have shown that recombinant dsRNA can be produced in the chloroplast of the edible microalga, *Chlamydomonas reinhardtii* and used to control disease in shrimp. Here we report a significant improvement in antiviral dsRNA production and its use to protect shrimp against white spot syndrome virus (WSSV). A new strategy for dsRNA synthesis was developed that uses two convergent copies of the endogenous *rrnS* promoter to drive high level transcription of both strands of the WSSV gene element in the chloroplast. New vectors were designed that allow rapid Golden Gate-mediated assembly of a transformation plasmid in which the ‘dual promoter-WSSV DNA’ cassette is targeted into the chloroplast genome, with selection based on the restoration of photosynthesis. PCR analysis of transformant lines confirmed the integration of the cassette and homoplasmy of the polyploid genome. Transcribed sense and antisense VP28-RNA were hypothesised to form an RNA duplex in the chloroplast stroma, and quantitative RT-PCR indicated that ∼100 μg dsRNA is produced per litre of transgenic microalgae culture. This represents an ∼10,000-fold increase in dsRNA relative to previous reports using convergent *psaA* promoters. The engineered alga was assessed for its ability to prevent WSSV infection when fed to shrimp larvae prior to a challenge with the virus. Survival of shrimp fed with dsRNA-expressing *C. reinhardtii* was significantly enhanced (68.3%) relative to the negative control. The study suggests that this new dsRNA production platform is significantly more efficient than that reported previously, and merits further scale-up and downstream processing studies.

## 1. Introduction

White Spot Syndrome Virus (WSSV) is the causative agent of White Spot Syndrome Disease (WSSD) in penaeid shrimp, including commercially relevant species such as Black Tiger prawns, King Prawns, and Atlantic White Shrimp. WSSD is widely regarded as the most detrimental disease seen in commercial aquaculture of these organisms, causing up to 100% mortality within 3-5 days (Oakey et al. 2019). The virus was first identified during a 1992 outbreak in mainland China and Taiwan, where it is still endemic in shrimp cultivation.

The detrimental commercial impact of WSSV has led to considerable research efforts towards developing mitigation strategies, with several biotechnological measures reported to decrease and delay losses in shrimp populations under lab-scale conditions (Flegel et al., 2008). Suitable antiviral agents are significantly limited compared to the wide range of antibiotics that target bacterial pathogens. However, RNA interference (RNAi) strategies have been reported to give a high degree of protection against WSSV infection. Delivery of double-stranded (ds) RNA specific to WSSV sequences into the shrimp cell triggers the RNAi mechanism, which consequently degrades invading viral RNA, thus halting the infection (Krishnan et al. 2009). A key mediator of WSSV entry into the host cell is the major envelope protein VP28, making it an attractive target for RNAi. Specific silencing of the *vp28* gene has been shown to reduce viral copy number in infected shrimp, resulting in a reduction in shrimp mortality (Xu et al. 2007), and the feeding of *Escherichia coli* expressing *vp28* dsRNA to shrimp yielded survival rates up to 80% relative to untreated shrimp (Thammasorn et al. 2015). Although promising, this strategy does not lend itself to commercial application, not lease due to concern over the dietary effects of *E. coli* on the shrimp, and the potential for the release of antibiotic resistance genes into the environment.

Microalgae are a broad class of photosynthetic unicellular eukaryotes, and are found in fresh, salt, and brackish waters. There has been a significant rise in interest in microalgae as feed and food ingredients over the past two decades, largely due to their sustainable nature and rich nutritional profile. Examples of commercially available microalgae for human consumption include *Chlorella vulgaris, Dunaliella salina, Haematococcus pluvialis*, and *Euglena gracilis*, with many other species forming an important component of animal feeds, including for shrimp and fish larvae (Muller-Feuga 2000). Genetic manipulation of the nuclear genome is possible for a considerable number of microalgae species, with a smaller subset also amenable to chloroplast genome modification (Spicer and Purton 2016). One application of these technologies is the production of functional biomolecules for oral delivery to aquatic animals (Fajardo et al. 2020).

*Chlamydomonas reinhardtii*, a freshwater microalga, is arguably the most studied microalgal species, and as such has a considerable set of molecular tools (Mussgnug 2015). These include well annotated nuclear and chloroplast genomes, established transformation protocols, and substantial parts libraries for gene expression. *C. reinhardtii* is able to grow photosynthetically, and also heterotrophically if provided with a suitable fixed carbon source. The ability to dispense completely with photosynthesis has made it a model organism for studying photosynthesis, but also allows the restoration of light-driven growth to be used as a transformation selection marker, side-stepping traditional markers based on antibiotic resistance genes (Economou et al. 2014). Although *C. reinhardtii* does not provide specific nutritional benefits such as the long chain PUFAs seen in many marine microalgae, its GRAS (Generally Recognised As Safe), status and ease of genetic manipulation makes it an attractive chassis for production and oral delivery of biomolecules (Rosales-Mendoza et al. 2020). We previously reported the first case of *C. reinhardtii* as an expression platform for dsRNA targeting of yellow head virus, and demonstrated partial protection against the virus in *Penaeus vannamei* shrimp (Charoonnart et al. 2019). The lack of full protection is thought to be due to the relatively low yield of antiviral dsRNA (16 ng/L of transgenic *C. reinhardtii* culture). The present study set out to remedy this shortfall by increasing the level of dsRNA produced in the *C. reinhardtii* chloroplast using a novel expression cassette, and to demonstrate the efficacy of the improved platform both by direct quantification of dsRNA, and by *in vivo* challenge of treated shrimp with WSSV.

## 2. Materials and Methods

### 2.1 Algal strain and maintenance

The *C. reinhardtii* strain CC-5168, also known as TN72, (*cw15*, Δ*psbH*) was used as a recipient line (Wannathong et al. 2016). The strain was maintained on Tris Acetate Phosphate (TAP) medium (Kropat et al. 2011) solidified with 2% agar, and grown under continuous low-intensity light (<10 μmol photon m^-2^ s^-1^) at 25 °C. Sub-culturing was performed monthly. Two full loops of healthy *C. reinhardtii* from solid medium were used to inoculate 400 ml TAP for liquid cultivation.

### 2.2 Plasmid design and construction

The p2xTRBL dsRNA expressing cassette was constructed using a Golden Gate approach modeled on the STEP platform (Taunt et al. manuscript in press). Three level 0 plasmids were built containing respectively: the 5’ *atpB*_Terminator/*rrnS*_Promoter element, the mRFP cassette, and the 3’ *rrnS*_Promoter/*rbcL*_Terminator element. The *C. reinhardtii* expression elements were amplified from genomic chloroplast DNA, with the addition of adaptor sequences to allow scar-free fusion of the promoter/ terminator pair, and insertion into the Level 0_Empty recipient vector. Also included were additional adapter sequences to allow the three Level 0 constructs to be assembled into the final Level 1 vector, p2xTRBL. The mRFP cassette was synthesized *de novo*, incorporating both adapter sequences for assembly, and additional sequences to allow dsRNA target sequences to be cloned into the final vector. Construction of this vector is summarized in Supplementary Figure 1.

A 307 bp fragment of the WSSV VP28 gene was amplified by PCR from a crude virus preparation with the primers designed to include inwards facing Esp3I sites and specific fusion sites for cloning into p2xTRBL (Supplementary Table 1). The Golden Gate reaction was conducted with 1x CutSmart buffer, 1 mM ATP, 400 units T4 ligase, and 10 units Esp3I at plasmid to fragment molar ratio of 1:8. The reactions were incubated at 37°C for 15 min, followed by 30 cycles of 16/37°C (2 min each), with a final digestion step at 37°C for 15 min, and heat inactivation at 65°C for 20 min. The resultant products transformed into DH5α and plated onto LB+Tet^10^ agar. White colonies were picked and the correct insertion of the WSSV fragment confirmed by PCR. The resulting plasmid was named p2xTRBL_VP28. The dsRNA expression cassette from this plasmid was then sub-cloned into the *C. reinhardtii* chloroplast transformation vector pSS116 following a similar protocol but using the dual outward facing BsaI sites, and with transformants plated on LB+Amp^100^ (Figure 1). White colonies were again confirmed by PCR, with the final construct named pSS116_VP28. Additional confirmation of correct cloning was achieved by diagnostic restriction endonuclease digestion using XhoI and BstXI. This plasmid was further amplified in *E. coli* DH5α for *C. reinhardtii* transformation.

**Figure 1.**
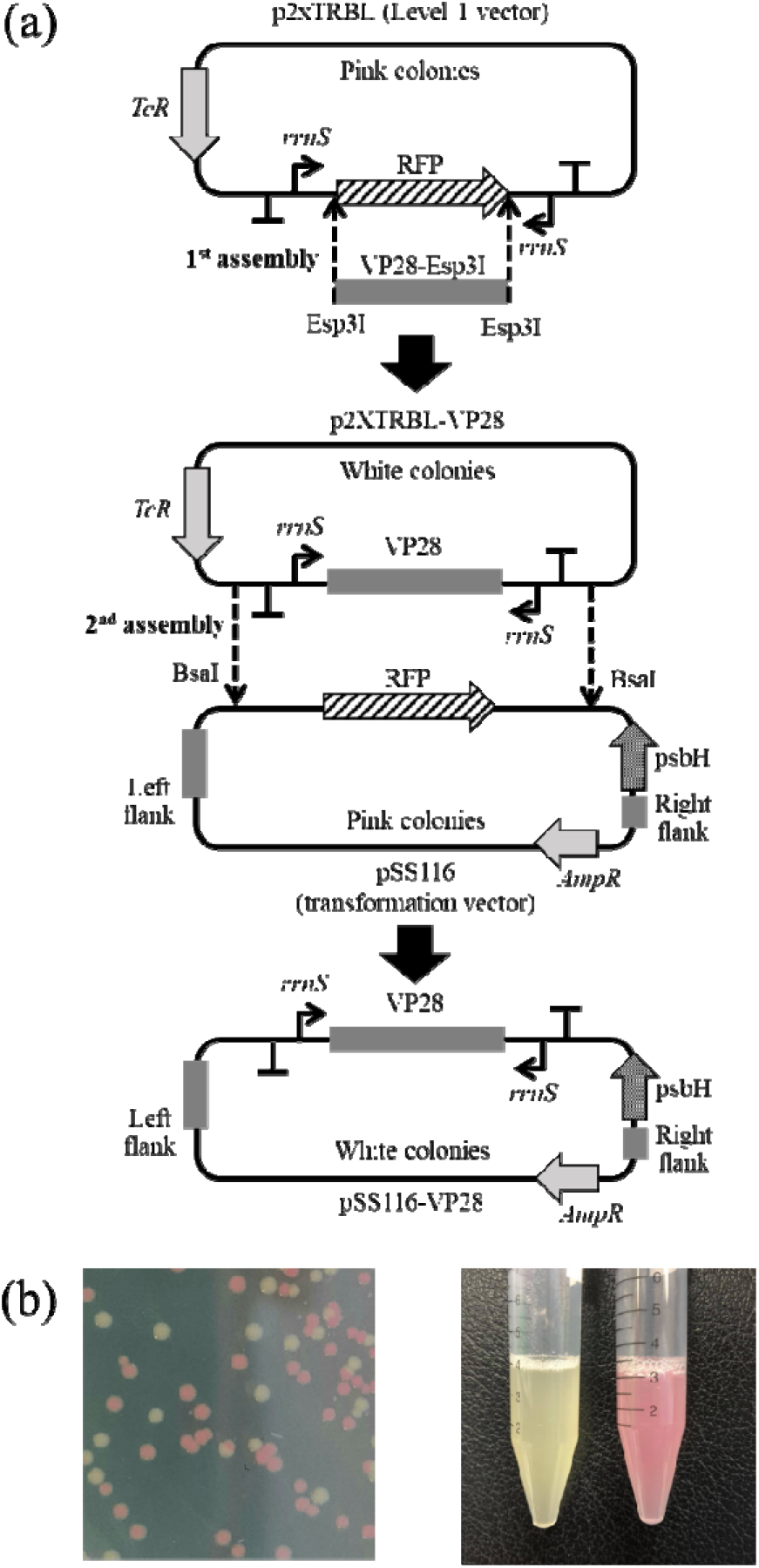
Schematic diagram for the construction of the dsRNA transformation plasmid pSS116-VP28. (a) Construction involved a two-step cloning strategy in which the VP28 DNA was first cloned into p2xTRBL using the type IIS restriction enzyme Esp3I and then the dual transcription cassette was cloned into pSS116 using a second type IIS enzyme, BsaI to flank the cassette with chloroplast DNA elements for recombination into the chloroplast genome. (b) Cloning was aided at each stage by a simple ‘pink/white’ screening system where transformant colonies and cultures carrying the correctly assembled plasmid gave rise to a ‘white’ phenotype.

### 2.3 Confirmation of dsRNA expression in E. coli

pSS116-VP28 was transformed into the RNase III deficient *E. coli* strain HT115(DE3) (Papić et al. 2018) and colonies selected on LB agar containing 100 μg/ml ampicillin. A single colony was inoculated into liquid LB+Amp^100^ broth and continuously shaken at 37°C for 18 hours. Re-inoculation was performed at 1:100 into fresh LB+Amp^100^ broth and continuously shaken for 8 hours following by cell collection by centrifugation. Total RNA was extracted using the phenol-chloroform (1:1) extraction method, and dsRNA purified according to Saksmerprome et al. (2009). Purified dsRNA was visualized by agarose gel electrophoresis and kept at -20 °C for further use as a standard for detection of dsRNA from transformant microalgae.

### 2.4 Chloroplast transformation and transformant selection

Chloroplast transformation of the cell wall-deficient strain TN72 was performed using the glass bead transformation method (Economou et al. 2014). 10 ml of starter culture was inoculated into 400 ml TAP medium and cultured with continuous shaking under low-intensity light until late log-phase (OD_750_ at 1.0–1.2). Then, cells were concentrated 100-fold to obtain a final cell density at approximately 2×10^8^ cells/ml. 300 μl of concentrated cells were aliquoted into 5 ml tubes containing 300 mg pre-sterilized 400-625 μm glass beads along with 5 μg of pSS116-VP2, and vortexed at maximum speed for 15s. High Salt Minimal (HSM) medium supplemented with 0.5% agar was melted and cooled to 42 °C prior to addition to the mixture followed by immediate pouring onto HSM agar plates. The transformation plates were incubated overnight in the dark cultured under 50 μmol photon m^-2^ s^-1^ white light at 25 °C. Successful transformation colonies where photosynthesis had been restored were obtained after 4–6 weeks.

Single colonies were individually picked and streaked on fresh HSM plates and continuously cultured for 3-4 weeks before next passaging. Homoplasmy was monitored by PCR following extraction via a modified CTAB method from Juntadech et al. (2012). To assure chloroplast DNA extraction, Rubisco-Large Subunit primers (RbcL_F and RbcL_R) were used as an internal control for PCR analysis (Charoonnart et al. 2019) (Supplementary Table 1). The VP28 fragment was detected using specific primers, VP28_ESP_F and VP28_ESP_R, to confirm insertion of the convergent *rrnS* promoter cassette. The primers TN72_F and TN72_R were used to confirm homoplasmy of the transformed chloroplast genome. Once confirmed to be homoplasmic, the transformant line was named TN72-dsVP28, and was further cultured for dsRNA quantification and viral protection studies. The pSS116 plasmid was also transformed using the same method to generate a photosynthesis-positive TN72 strain, called TN72-SS, that was used as a negative control for further experiments.

### 2.5 Double-stranded RNA detection and quantification

A full loop of TN72-dsVP28 was inoculated into 400 ml TAP medium and cultured by shaking at 100 rpm under continuous light (50 μmol photon m^-2^ s^-1^) at 25 °C. The culture was harvested (centrifugation at 3,000 x *g* for 5 min), at late log phase (day 4) and the 1^st^ day of stationary phase (day 5) (growth analysis data not shown) and re-suspended with Trizol solution (Ambion, USA) for RNA extraction to obtain total RNA. Undesired DNA and single-stranded RNA (ssRNA) were removed by DNase I and RNase A treatment as per the manufacturer’s instructions to obtain isolated dsRNA. VP28 specific dsRNA was detected by RT-PCR (RBC Bioscience, Taiwan) using VP28_ESP_F and VP28_ESP_R primers. The formation of dsRNA was confirmed by observing degradation of dsRNA by RNase III treatment.

Quantitative RT-PCR (qRT-PCR) was performed using the same set of primers as for RT-PCR (KAPA Biosystem, USA). Isolated dsRNA was incubated at 95 °C for 5 min and immediately chilled on ice for 2 min prior to addition into the qRT-PCR reaction to ensure denaturation of dsRNA. A series of ten-fold dilution of purified dsRNA from the bacterial expression was used to generate a standard curve. The amount of standard dsRNA was calculated according to a formula obtained from the standard curve (R^2^=0.958).

### 2.6 Hanging-bag culture and biomass preparation

Pre-inoculum of TN72-SR, a photosynthesis restored strain from Charoonnart et al. (2019), and TN72-dsVP28 was prepared by inoculating a full loop of the strain grown on TAP-agar plates in 100 ml TAP medium and shaking under continuous light at 25°C for 3-4 days. Absorbance at 750 nm was measured and calculated for a volume input into the polythene tubular bag containing 17 L TAP medium to make a starter absorbance at 0.1. The hanging bag photobioreactor was set up according to Cui et al. (2021). Filter-sterilized air was provided from the bottom of the bag and continuous illumination provided at 100 μE m^-2^s^-1^ by Osram Lumilux Cool daylight fluorescence tubes. The bag cultures were cultivated for 5 days at 25°C with growth monitored by measuring absorbance at 750 nm every 12 hours. At harvesting time, chemical flocculation by FeCl_3_ at a final concentration 2 mM was performed with continuously air supplied for 10-15 min before leaving the bag to stand until the algae had completely precipitated. Microalgal biomass were recovered by centrifugation of the concentrated culture at the bottom of the bag at 4,500 rpm for 20 min and the paste was immediately subjected to freezing at -20°C. This was then lyophilized in a freeze-dryer over a period of 24 h, and the dried biomass was kept at -20°C until further analysis. Double-stranded RNA purification and detection were performed for confirmation of the accumulation of specific dsRNA, as above.

### 2.7 In vivo protection assays of VP28-dsRNA expressing C. reinhardtii against WSSV in shrimp

#### 2.7.1 Preparation of animal and viral inoculum

Specific Pathogen Free (SPF) Post larval (PL) (23 day old) *Penaeus vannamei* shrimp were purchased from a major hatchery farm (supplier wishes to remain anonymous) and acclimatized in 10 parts per trillion (ppt) salinity artificial seawater for 7 days prior to use as an animal model. Average animal mass and length were approximately 0.03 g and 1.5 cm respectively on the first day of experiment. To prepare the WSSV inoculum for the challenge assay, a mixed gender group of 20 shrimps (10-15 g each) were fed with WSSV infected homogenized shrimp tissue to a rate of 10% of total body mass. Visibly infected shrimps were euthanized and muscle and pleopod tissue collected. WSSV infection was confirmed by the IQ2000 PCR-based detection kit (GeneReach Biotechnology Corp., Taiwan). Infected shrimp muscle tissue was homogenized and kept at -20 °C for later use in the oral challenge assay.

#### 2.7.2 Assay for viral protection in shrimp

The experiment was dividing into four feeding groups of 100, representing four replicates of 25 animals. Each replicate was cultured in its own glass tank containing 2 L of 15 ppt artificial seawater with continuous aeration. Groups one and two represented negative and positive controls respectively and were fed with commercial feed twice daily at 5% body weight throughout the experiment. Groups three and four were fed with commercial feed supplemented at 1:1 ratio of TN72-SR (negative control transformant line from Charoonnart et al. (2019)) and TN72-dsVP28, respectively, at the same amount as group one and two. The WSSV challenge was conducted on day four of the experiment. Groups two, three, and four were exposed to oral administration of WSSV infected tissue at a rate of 50% total body mass (1.25×10^8^ WSSV copies/tank). Group one was not exposed. Animal survival was observed daily until group 2 reached 100% mortality.

## 3. Results

### 3.1 Design of the p2xTRBL dsRNA expression vector

The p2xTRBL vector is based on the principle of convergent promoter elements, each facing an in-sense terminator located on the opposite side of a Golden Gate cloning region, and downstream of the opposite facing promoter (Figure 1A). For this vector, the promoter of the *Chlamydomonas reinhardtii* chloroplast gene *rrnS*, which encodes the 16S ribosomal RNA, was chosen as it is the most transcriptionally active promoter in the *C. reinhardtii* chloroplast (Blowers et al. 1990). Insertion of the target sequence is mediated by Golden Gate cloning using the type IIS restriction enzyme Esp3I as discussed below. Once assembled, the expression cassette comprising the target sequence flanked by the promoters and terminators can be excised using a second type IIS enzyme, BsaI and transferred to a chloroplast integration vector such as pSS116 through a second Golden Gate cloning step, as illustrated in Figure 1A. The presence of different antibiotic selection markers on the two vectors avoids the need to purify the excised cassette prior to cloning into pSS116.

### 3.2 Golden Gate assembly allows for quick and highly efficient construction of plasmids

The insertion of the *VP28* PCR product into the p2xTRBL vector, and of the resulting expression cassette into the pSS116 vector, were both conducted by one-pot-one-step Golden Gate reactions. This method allows for the directional cloning of one or more fragments into a vector without having to separate digestion and ligation steps: the use of type IIS restriction endonuclease (RE) sites ensure that once the desired plasmid is assembled the enzyme recognition sites are lost, precluding it from any further digestion. Both vectors also featured a colorimetric negative screening device, such that successfully assembled plasmids lose the bacterial mRFP expression cassette located between the pair of RE sites and thus yield white colonies, as opposed to the pink colonies generated by the parental plasmids (Figure 1B). Using a 1:8 molar ratio of p2xTRBL to *VP28* DNA we achieved 90% successful assembly based on colony colour. All selected white colonies (26) were shown by PCR analysis to be correctly assembled p2xTRBL-VP28.

Downstream cloning of the expression cassette into pSS116 was conducted using a similar method, but with BsaI used as the restriction endonuclease instead of Esp3I, and colonies selected on ampicillin as opposed to tetracycline. Assembly as judged by pink/ white selection was at a similarly high level (98% white colonies), and again all picked colonies gave the expected PCR products for the recombinant plasmid, pSS116_VP28. A diagnostic digestion of the purified plasmids using XbaI and BstXI further confirmed this.

### 3.3 The dual-rrnS promoter system is capable of dsRNA production in E. coli

After assembly of the pSS116_VP28 plasmid in the standard *E. coli* strain DH5α, the plasmid was transformed into the RNase III deficient strain HT115(DE3) to allow accumulation of dsRNA (Papić et al. 2018). Cultures containing either pSS116_VP28 or the recipient pSS116 were grown to mid-log phase (8 hours), and the purified dsRNA analysed by agarose gel electrophoresis. As shown in Figure 2a, a dsRNA band of the expected size of ∼0.4 kb is seen for the *VP28* dsRNA from the pSS116_VP28 culture, whereas no such band was observed from the pSS116 culture. Total dsVP28 accumulation was estimated from band intensity to be approximate 500 μg. Furthermore, RT-PCR using specific primers to the *VP28* sequence was conducted to confirm the presence of VP28 dsRNA. An amplicon of ∼0.3 kb representing just the VP28 sequence without the upstream and downstream transcribed regions was observed from pSS116-VP28, while this amplicon was not seen for the pSS116 line (Fig. 2b).

**Figure 2.**
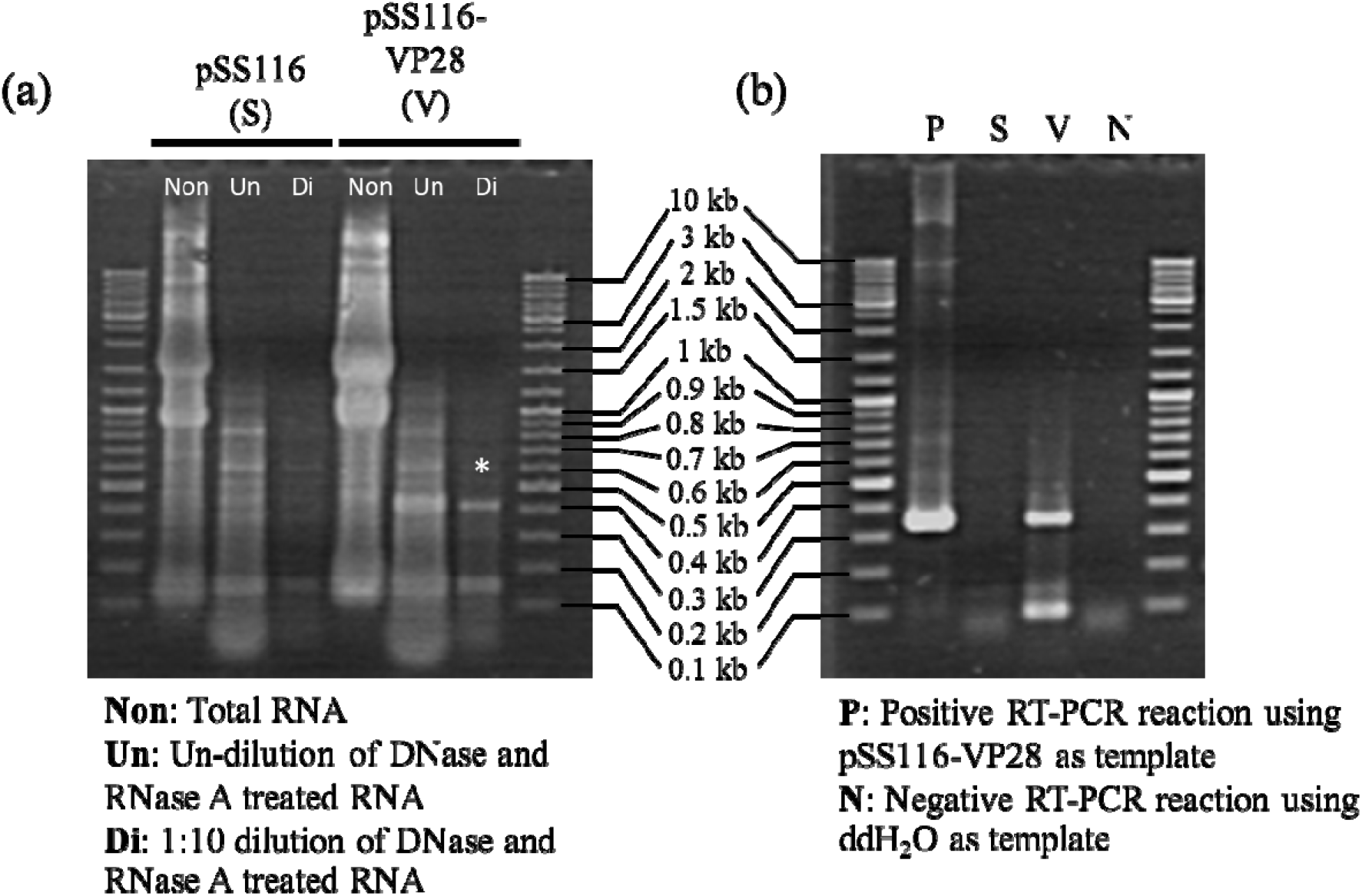
Purified RNA from *E. coli* HT115(DE3) transformed with pSS116 (S) and pSS116-VP28 (V), showing total RNA (Non), then presumably RNA digested with DNase and RNase A (Un) and 1:10 diluted and treated RNA (Di) (a). RT-PCR of DNase and RNase A treated RNA with dsVP28 specific primers (b). Lane N and P represent negative and positive reaction for PCR analysis using ddH_2_O and dsVP28, respectively, from bacterial system as a template.

### 3.4 Generation of C. reinhardtii transplastomic lines

The pSS116 chloroplast integration vector contains *C. reinhardtii* chloroplast sequences flanking the cloning site such that the cloned DNA is targeted into the genome at a neutral site between *psbH* and *trnE2*. Furthermore, a wild-type copy of *psbH* on the right flank serves as a selectable marker, rescuing the Δ*psbH* mutant TN72 to phototrophy (Wannathong et al. 2016). Chloroplast transformants of TN72 were generated using pSS116-VP28, and pSS116 as a control, with the first colonies becoming visible on minimal medium after six weeks. A total of seven colonies were observed for pSS116-VP28 and only two for pSS116 transformation.

After the second re-streak on minimal medium, DNA was extracted and PCR analysis conducted, with amplification of the endogenous gene *rbcL* used as a positive control for extraction (Fig. 3a). The *VP28* DNA was successfully amplified from the pSS116_vp28 lines, and not seen from the pSS116 lines (Fig. 3b). As the copy number of the chloroplast genome in *C. reinhardtii* cells is typically ∼80 (Gallaher et al. 2018), and transformation initially results in a heteroplasmic state with both transformed and original copies of the genome within the cell (Purton 2007), then homoplasmy in the transformant lines was assessed using primers specific for the original TN72 genome. The absence of a band from the pSS116_VP28 and pSS116 lines suggesting homoplasmy (Fig. 3c).

**Figure 3.**
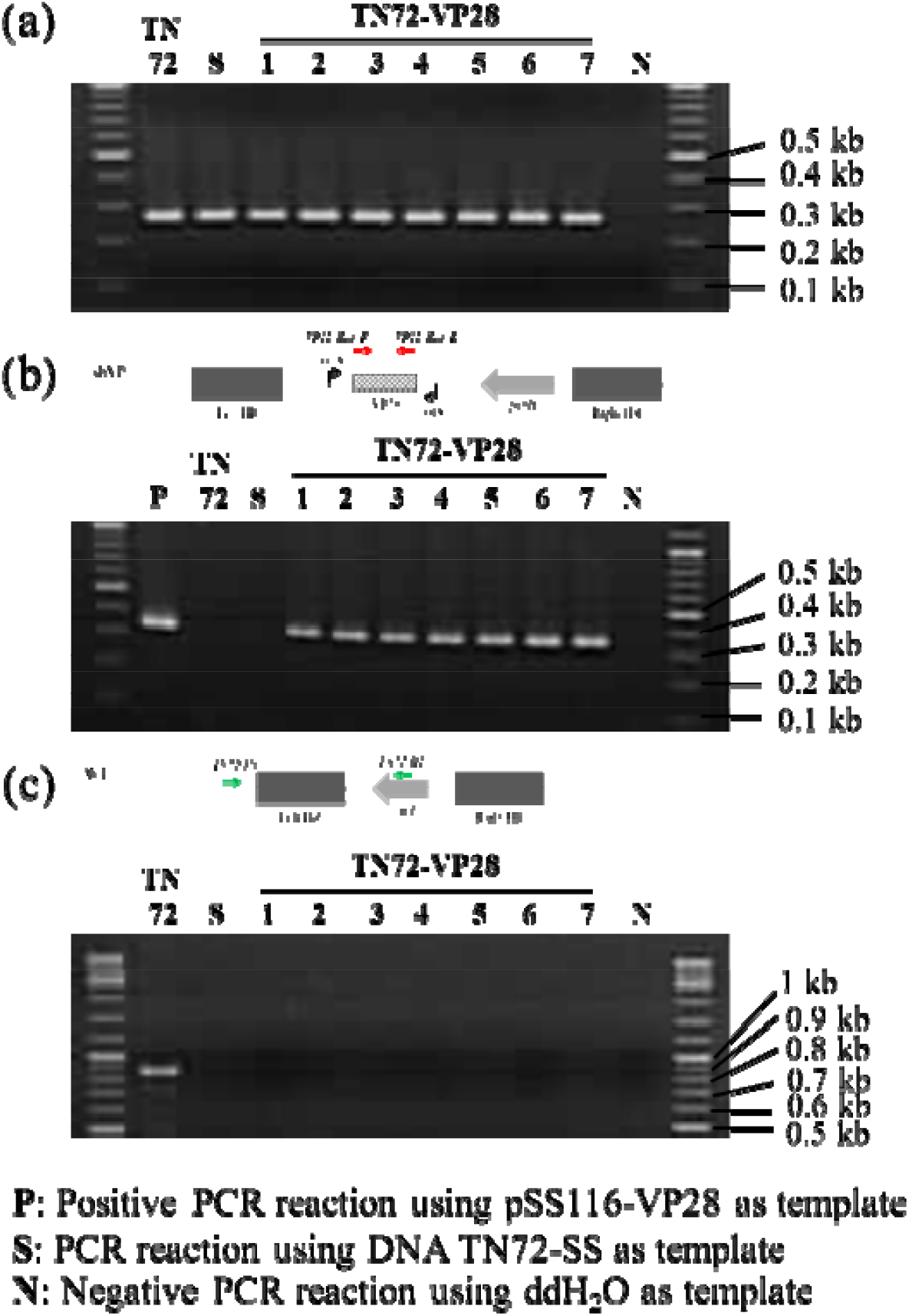
PCR analysis for selection of homoplasmic transgenic microalgae (TN72-VP281-7) containing the dsRNA expressing cassette. Primers rbcL_F and rbcL_R for chloroplast DNA control comparing to TN72 and original pSS116 transformed strain (S) (a), schematic of insertion region displaying VP28_Esp_F and VP28_Esp_R binding for VP28 fragment detection (b), and TN72_F and TN72_R for remaining plastid detection. Lane N and P represent negative and positive reaction for PCR analysis using ddH_2_O and dsVP28, respectively, from bacterial system as a template.

One representative transformant line containing the 2xTRBL_VP28 cassette was selected for dsRNA investigation and named TN72-dsVP28, with a corresponding pSS116 line chosen as a negative control and named TN72-SS.

### 3.5 Convergent psaA promoters significantly enhanced production VP28-dsRNA relative to the previous study

The yield of dsRNA in the TN72-dsVP28 strain was analysed following total RNA extraction, and DNase I and RNase A treatment to digest all DNA and ssRNA leaving only dsRNA, as previously demonstrated in Charoonnart et al. (2019). The specific presence of VP28 dsRNA in the TN72-dsVP28 RNA sample, but not in an equivalent sample from the TN72-SS control strain, was investigated by RT-PCR. TN72-dsVP28 gave the expected amplicon at 307 bp, whereas no band was seen for TN72-SS (Fig. 4a). Furthermore, this amplicon was lost when the dsRNA template was treated with the dsRNA specific enzyme, RNase III (Fig. 4b), confirming that VP28 dsRNA was indeed present in the TN72_VP28 line.

**Figure 4.**
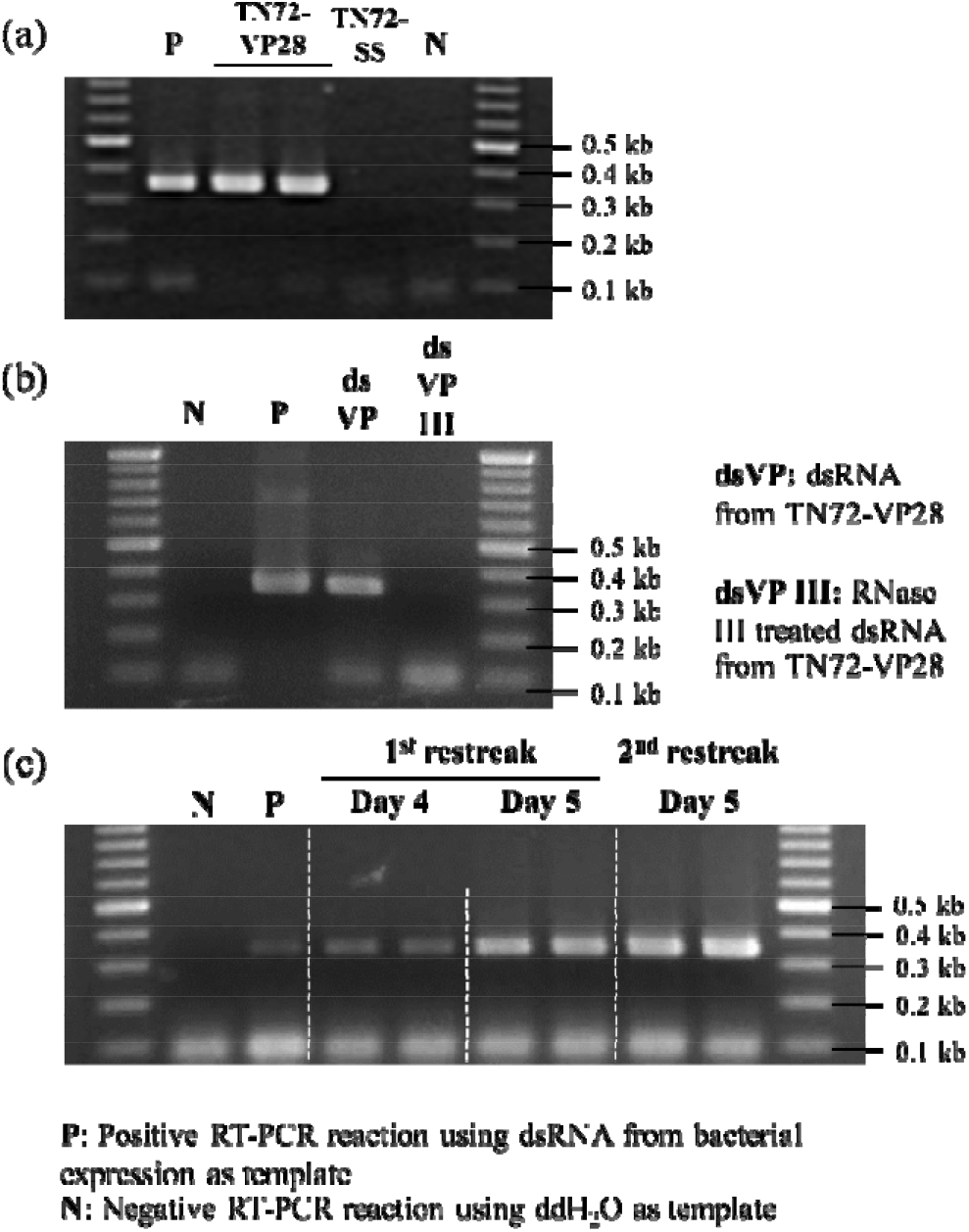
RT-PCR analysis for detection of dsRNA from dsRNA-expressing cassette containing strain; (a) dsRNA purified from TN72-VP28 shows the VP28 amplicon, whereas TN72-SS strain does not, (b) TN72_VP28 dsRNA does not show the VP28 amplicon after RNase III treatment, and (c) the determination of dsRNA yield at different harvesting time. Lane N and P represent negative and positive reaction for PCR analysis using ddH_2_O and dsVP28, respectively, from bacterial system as a template.

To investigate whether growth phase had an effect on dsRNA expression, semi quantitative RT-PCR was performed on samples harvested after four and five days of culture. These data show significantly higher expression of dsRNA from the 5 day culture suggesting accumulation of dsRNA in early stationary phase (Fig. 4c). Fully quantitative qRT-PCR was then used on the sample harvested on the 5^th^ day to give a conclusive dsRNA yield. Melt curve analysis using purified dsRNA from the HT115(DE3)_VP28 bacterial expression system gave a melting temperature (Tm) of 82.40±0.32 °C while dsRNA obtained from TN72-dsVP28 strain gave a Tm of 82.48±0.16 °C. Quantitative analysis gave 41.13 pg VG28 dsRNA from a 500 ng dsRNA input, which corresponds to 119 μg of *VP28* dsRNA from 1 L of TN72-dsVP28 culture (OD_750_ at 3.9) while a previous expression cassette using the *psaA* promoter as described in Charoonnart et al. (2019) yielded 16 ng specific dsYHV from 1 L of culture (OD_750_ at 1.8).

### 3.6 Shrimp fed with the TN72-dsVP28 line show ∼70% survival against WSSV

To evaluate the efficiency of TN72-dsVP28 in protecting against WSSV, a feeding experiment following by a WSSV challenge assay was performed. Prior to supplementing the feed with transgenic microalgae, cell biomass of both TN72-SR and TN72-dsVP28 was analysed for the present of specific dsVP28 accumulation by RT-PCR analysis and the result confirmed the present of dsRNA in TN72-dsVP28 but not TN72-SR (Supplementary data). As shown in Figure 5, a lethal dose 50 was reached on days 4 or 5 post infection (dpi) in shrimp receiving the TN72-SR supplement and positive group (commercial feed without algae, full WSSV challenge), respectively whereas the negative group (commercial feed without algae, no WSSV challenge) showed 91-94% survival. In contrast, the shrimp that received the feed supplemented with TN72-dsVP28 showed a marked reduction in mortality compared to the TN72-NR control with 69.49±6.25% at 6 dpi which then remained at this level (Figure 5). On the last day of the experiment (8 dpi), the survival of both the positive and TN72-SR supplemented groups was less than 10%,whereas shrimp receiving feed supplemented with TN72-dsVP28 showed survival of ∼68%. Statistical analysis of these numbers gives significant at 99% confidentiality.

**Figure 5.**
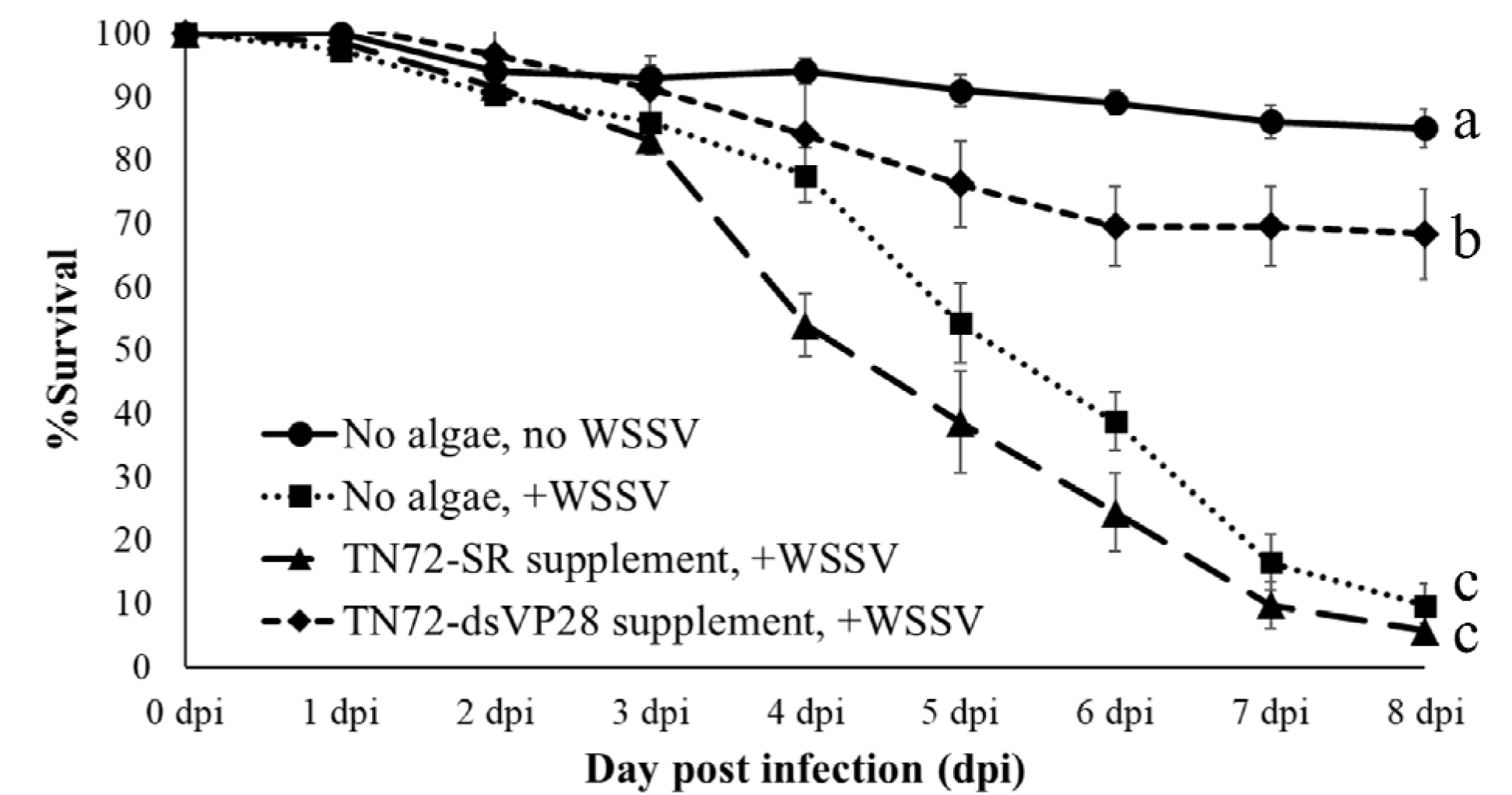
Percentage survival of shrimp during the feeding and challenge trial. Comparing between WSSV(-) (no WSSV challenge) and WSSV(+) (WSSV challenge), TN72-SR and TN72-VP28 supplement prior WSSV oral challenge. Bar represents standard error. Alphabet a, b, and c represent homogeneous subset by Duncan’s test of the percentage survival of shrimps from 8 dpi.

## 4. Discussion

The novel vector p2xTRBL and corresponding transformation vector pSS116 represent a significant improvement in the construction and efficacy of dsRNA expressing lines of *C. reinhardtii*. The use of convergent *rrnS* promoters marks a significant improvement over a previous design where *psaA* promoters were employed (Changko et al. 2020; Charoonnart et al. 2019; Gangl et al. 2015; Zedler et al. 2015). This is to be expected, as the *rrnS* promoter has been shown previously to be the most transcriptionally active promoter in the *C. reinhardtii* chloroplast (Rasala et al. 2011); however, it should also be noted that the p2xTRBL cassette represents the first fully purpose-built *C. reinhardtii* dsRNA expression system. Charoonnart et al. (2019) previously constructed a dsRNA expressing cassette by modifying pSRSapI to contain convergent *psaA* promoters, but as the original vector was designed for protein synthesis it not only contained unnecessary 5’UTR regions, but also only the sense promoter had a corresponding transcriptional terminator. The antisense promoter on the other hand, will drive transcription until a suitable native terminator is reached, lending the dsRNA an extended ssRNA “tail” as well as wasting cellular resources.

Since chloroplast promoters are prokaryotic in origin, they are often active in *E. coli*, with the *rrnS* promoter being no exception. This presented a useful opportunity to validate the dsRNA-expressing cassette in pSS116_VP28, prior to algal transformation, and also to produce a stock of VP28 dsRNA to be used as a positive control. Building on several reports where convergent bacteriophage T7 promoters have been used to produce dsRNA in *E. coli* (García et al. 2015; Kim et al. 2015), we transformed the pSS116_VP28 vector into the RNase III deficient *E. coli* strain HT115(DE3). The resulting system was shown to be able to produce reasonable amount of dsRNA, but the amount was lower than that produced from hairpin expressing cassettes (Chen et al. 2018). Nonetheless, hairpin cassettes are not stable in plastids due to the presence of a RecA recombinase that might loop out the inverted repeat in hairpin cassette (Nakazato et al. 2003). A convergent cassette was therefore considered preferable for dsRNA production in the *C. reinhardtii* chloroplast.

The use of traditional antibiotic resistance selection markers for engineering of microalgae to be used in aquaculture is highly undesirable because of the risk of spreading such cassettes to other microorganisms by horizontal gene transfer. The ability for chloroplast transformants to be selected without such markers is hence a large advantage, as has be demonstrated numerous times (Changko et al. 2020; Charoonnart et al. 2019; Gangl et al. 2015; Zedler et al. 2015). Despite these benefits, the transformation with pSS116_VP28 suffered from long incubation times and low transformation efficiency. The incubation period was 2 weeks longer than previous *psaA* convergent promoter cassette (4 weeks), and those reported for protein production (Gangl et al. 2015). The delay of transformation and low efficiency may have been due to the presence of the convergent *rrnS* promoters, with the inverted repeat potentially interfering with homologous recombination; however, as colonies were ultimately recovered and shown to all be correct. Moreover, the transformant strains rapidly reached homoplasmy, showing no evidence of the recipient genotype after only two rounds of single-colony isolation. Similar results have been achieved recently for two transformant lines where the *rrnS* convergent promoter system has been used to produce dsRNA targeting the RdRp gene of YHV and VP9 and ORF366 of WSSV (unpublished results).

The resultant transplastomic line, TN72-VP28, was shown to express VP28-specific dsRNA by RT-PCR following selective degradation of DNA and ssRNA. It was then not possible to amplify the VP28 sequence following the use of RNAse III to specifically degrade dsRNA. Combined, these two results give strong evidence for the production of VP28 dsRNA in the *C. reinhardtii* chloroplast. Subsequent RT-qPCR was used to quantify the VP28 dsRNA, giving a yield of 119 μg of VP28 dsRNA from 1 L culture at an OD_750_ of 3.9. Though the dsRNA yield obtained is very much lower than a bacterial system, the value of the algal system is that it can be used as a whole-cell feed. Furthermore, the marker-less transformation method means that the only transgenic DNA in the feed is the ∼0.3 kb section of the viral VP28 gene, and this DNA is endemic anyway in an infected shrimp pond. This current study representing a ∼10,000-fold increase in dsRNA compared to the earlier work using the *psaA* promoter, although as this work expressed a different target sequence it is not possible to make a direct comparison. It is of note that levels of dsRNA were observed to increase from day four to day five of the cultivation period. Further work will be needed to confirm this trend, and give a more complete picture of dsRNA expression over a full growth period. It would also be of interest to investigate whether other abiotic factors can influence dsRNA accumulation, for example the effect of light levels or media composition.

The drying of the algal biomass represents an effective method of killing the microalgae thereby reducing the chance of environmental contamination with a GM species when deployed in aquaculture. Furthermore, the drying produces a powder suitable for formulation into shrimp feed where the dsRNA is potentially stable at room temperature for extended periods. Further analysis is required to establish the exact half-life of the dsRNA under these conditions, together with the dose level needed for effective protection so that a full technoeconomic assessment can be made. Fasaei et al., (2018) reported that harvesting and dewatering are the key cost determinants in algal biomass production. The results revealed that feed supplemented with TN72-dsVP28 powder had the highest survival percentage compared to the group treated with TN72-SR (no dsRNA expression) at the last day of experiment. It could be implied that better protection are the results of dsRNA-mediated defence. The result is in accordance with the increasing of dsRNA comparing to Charoonnart et al. (2019) will confer to the improvement of *in vivo* protection efficiency. Enhancement of dsRNA expression level in algal system should be performed to obtain a minimum use of the powder for viral protection in the pilot and farm scale.

The *Penaeus vannamei* shrimp is known to be highly susceptible to WSSV throughout its life cycle, but the post larval (PL) stage of development is considered to be when it is must vulnerable to viral attack according to data collection from hatchery farms and nursery farms throughout Thailand by the Department of Fisheries (Kongkumnerd and Saleetid, 2019). It is therefore essential to ensure PL stocks are healthy and disease-free before transferring them to grow-out ponds. Since WSSV can be transmitted horizontally, viral transmission in the TN72-SR group was potentially quicker than in the TN72-VP28 group resulting in 100% mortality being reached more rapidly. Feeding PL shrimp with inactivated microalgae expressing dsRNA delayed the massive mortality of the shrimp following viral infection. Together with good farm management, shrimp farmers may therefore be able to rescue shrimp from mortality and gain some profits instead of completely discarding the whole crop..

## 5. Acknowledgement

This work was supported by Royal Society together with Thailand Science Research and Innovation (TSRI) through a Newton Advanced Fellowship to VS in collaboration with CR. Additional support was received from a Royal Society International Collaboration Award to CR and VS, an EU H2020 grant (‘PharmaFactory’ H2020 REA 774078) to SP, and Mahidol University (Fundamental Fund: Basic Research Fund: fiscal year 2022) (Grant no. BRF1-054/2565) to PC. The authors wish to thank Rebecca Moore for her assistance in the construction of p2xTRBL.

## 6. Statement of informed animal rights

Experiments related to shrimps used in this study were approved by BT-IACUC, protocol no. BT Animal 8/2562. All procedures performed in studies involving animal were in accordance with the guidelines of the Australian, New South Wales state government for the humane harvesting of fish and crustaceans.

