## Supplementary figure for "Transgenic microalgae expressing double-stranded RNA as potential feed supplements for controlling white spot syndrome in shrimp aquaculture"

### Enhanced production of antiviral dsRNA in the *Chlamydomonas reinhardtii* chloroplast via a novel convergent promoter expression system

Patai Charoonnart[1,2], Henry N Taunt[3], Conner Webb[4], Pantaree Limvatanyu[1], Saul Purton[3], Colin Robinson[4], and Vanvimon Saksmerprome[1,2]*

**Abstract**

The present work presents an improvement of microalgal antiviral dsRNA production for controlling disease in shrimp aquaculture. 307 bp of sequence targeting the VP28 gene of white spot syndrome virus (WSSV) was inserted between two convergent rrnS promoters in the novel vector p2XTRBL, which was then subcloned into the transformation vector pSS116 using Golden Gate assembly. The recombinant plasmid was transformed into the *Chlamydomonas reinhardtii* chloroplast, and transformants selected by the restoration of photosynthesis. The presence of the cassette and homoplasmy of the algal transformants was confirmed by PCR analysis. Transcribed sense and antisense VP28-RNA were hypothesised to form an RNA duplex in the chloroplast stroma, and quantitative RT-PCR indicated that ~100 µg dsRNA was obtained per litre of transgenic microalgae culture. This accumulation of dsRNA represents a 10,000-fold increase relative to previous reports using convergent *psaA* promoters. Recombinant *C. reinhardtii* was assessed for its ability to prevent WSSV infection in shrimp larvae by direct feeding. After WSSV challenge, the survival of shrimp treated with dsRNA-expressing *C. reinhardtii* was significantly enhanced (95.2%) relative to the negative control without dsRNA treatment. The study suggests that this new algal production platform for dsRNA is significantly more efficient than the previous report, and it merits further scale-up and downstream processing studies.

**Keywords**: *Chlamydomonas reinhardtii*; double-stranded RNA; shrimp diseases; white spot syndrome virus; chloroplast transformation

**Supplementary 1**

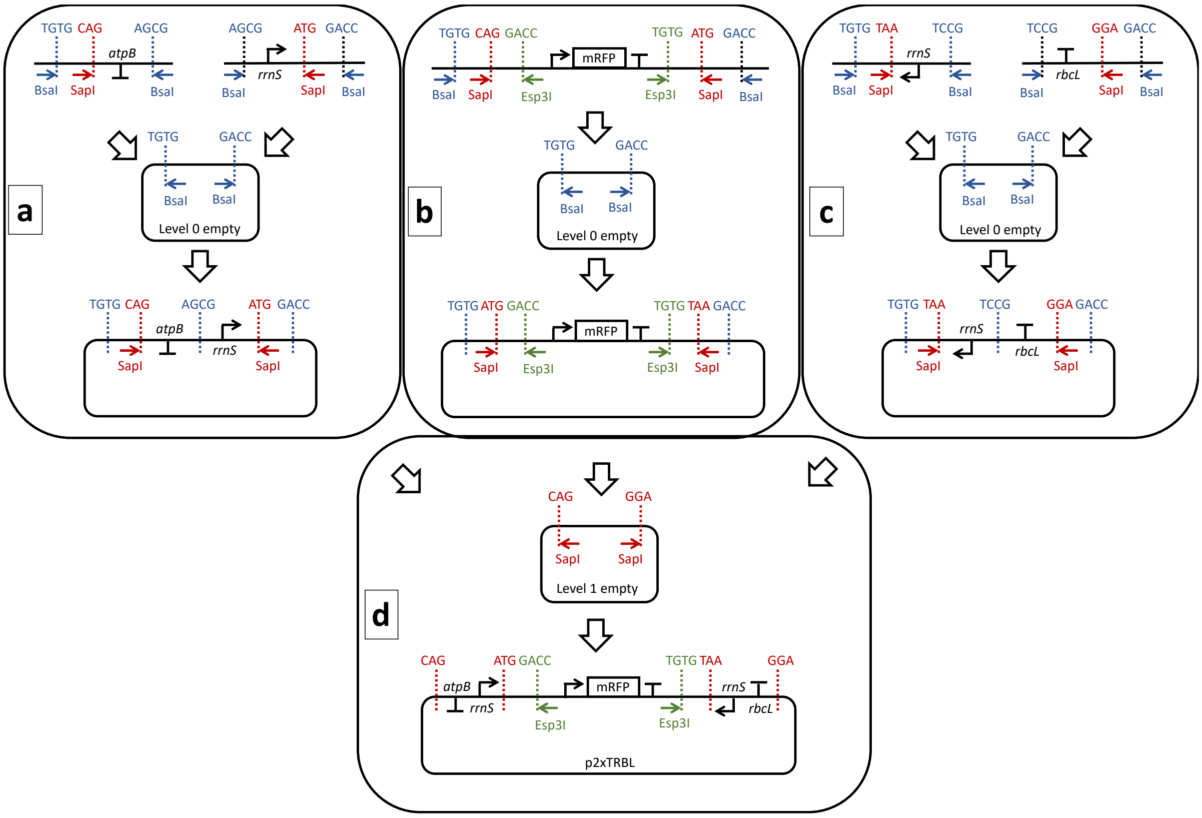


**figure 1.** *C. reinhardtii* expression elements were amplified by PCR, with the addition of 5' extensions to allow Golden gate cloning into the Level 0 empty vector (a, c). The mRFP cassette was synthesized *de novo*, also with the addition of Golden Gate cloning sequences (b). The three Level 0 cassette were cloned into a Level 2 empty vector to generate p2xTRBL (d).
